# From cellular synaptic input profiles to population-level respiratory circuitry

**DOI:** 10.64898/2026.11.23.768629

**Authors:** Yaroslav I. Molkov, Ilya A. Rybak, Jeffrey C. Smith

## Abstract

Breathing depends on coordinated activity among brainstem neurons. Their discharge patterns show when populations are active but do not specify the synaptic inputs that generate these patterns. Assuming that respiratory pattern generation involves a small number of interacting populations distinguished by discharge timing and excitatory or inhibitory output, we used synaptic conductances as readouts of their convergent activity. We reconstructed excitatory and inhibitory conductances from our previously described in situ rat brainstem-spinal cord recording archive (168 recordings, 150 neurons). We represented candidate presynaptic activity by five independently recorded population discharge patterns, assuming that phase-dependent inputs originated within these populations. We fitted each conductance time course as a nonnegative weighted sum of these patterns and tested whether candidate inputs could be omitted from the population mean, using variation across cells.

Post-inspiratory inhibition was supported in all three inspiratory populations.

Ramp-inspiratory neurons also showed augmenting-expiratory inhibition, and pre-inspiratory/inspiratory neurons showed augmenting-expiratory excitation. Excitation increased during the active phases of ramp-inspiratory and augmenting-expiratory neurons. Both expiratory populations showed inspiratory excitation alongside their prominent inspiratory inhibition. The inferred circuit retained major inhibitory interactions of the transection-based three-phase architecture and included additional excitatory inputs suggested by the earlier analysis. We distinguished some individual inspiratory discharge patterns but could not resolve other assignments. Source assignments depended on conductance reconstruction and how well the activity patterns represented presynaptic timing. These findings support excitatory interactions across populations with distinct discharge phases and suggest a contribution of late-expiratory excitation to recruitment of the next inspiration.

**Key points:**

- Respiratory discharge patterns show when neurons are active but not the synaptic interactions generating these patterns. Our recent work reconstructed previously unavailable excitatory and inhibitory synaptic inputs in respiratory neurons (Molkov et al., 2025).
- We reconstructed inputs in a larger sample from the same archive and combined them with published population activity to identify candidate input sources for each receiving population and reconstruct a circuit.
- The inferred circuit retained major inhibitory interactions of the three-phase architecture proposed to explain brainstem transection experiments. It included excitatory and inhibitory subpopulations with post-inspiratory and augmenting-expiratory discharge, with additional excitation during active and silent phases of receiving neurons.
- Some inputs could be assigned to individual inspiratory discharge patterns, assuming representative activity patterns; overlapping activity left other assignments unresolved.
- Excitatory inputs link populations active in different phases. Rising late-expiratory excitation together with declining inhibition provides a candidate mechanism for recruitment of the next inspiration.

## 1 Introduction

Each breath requires coordinated activation of inspiratory pump muscles, upper-airway muscles, and expiratory effectors. A brainstem respiratory network generates a necessary complex time-dependent output activity through interactions among rhythm-generating circuits, premotor pathways, and pontine and medullary inputs (Smith et al., 2009; 2013). The pre-Bötzinger complex (pre-BötC) contains a core inspiratory rhythm-generating circuit. Neuronal populations in the Bötzinger complex (BötC) and other regions of the ventral respiratory column in the medulla contribute to phase transitions and motor pattern formation. Defining how these populations are connected and interact in respiratory circuits remains a central problem in respiratory neurophysiology.

The three-phase pattern of the respiratory cycle comprises inspiration, post-inspiration (E1), and late expiration (E2). We analyze five neuronal discharge patterns expressed during these phases. Pre-inspiratory/inspiratory (pre-I/I) activity begins before inspiration and continues into inspiration. Early-inspiratory (early-I) activity is strongest near inspiratory onset, whereas ramp-inspiratory (ramp-I) activity increases during inspiration. Post-inspiratory (post-I) activity is strongest just after inspiration and declines during expiration. Augmenting-expiratory (aug-E) activity increases during E2. These categories describe discharge timing; each can include neurons with different transmitter identities, anatomical locations, and projections (Richter and Spyer, 2001; Rybak et al., 2007; Smith et al., 2007; 2013). Other documented respiratory discharge patterns are outside the present analysis.

Discharge patterns and their transformations after sequential brainstem transection have guided three-phase generating circuit concepts for several decades (Botros and Bruce, 1990; Balis et al., 1994; Rybak et al., 1997; 2007; Smith et al., 2007; Lindsey et al., 2012). A pontine-medullary in situ preparation expresses three-phase activity. Pontine removal produces a two-phase rhythm, and a further transection rostral to the pre-BötC leaves one-phase rhythmic inspiratory activity (Smith et al., 2007; 2009). These experiments establish when populations fire and which reduced networks remain rhythmic. They do not reveal which populations provide the excitatory and inhibitory synaptic inputs received by each discharge-pattern population. Circuit architectures with different connections can therefore remain compatible with the same recorded activity.

The recurring discharge patterns in cell and population recordings, together with their organized changes after transection, motivate a population-level description of respiratory circuitry (Smith et al., 2007; 2013). We adopt the working assumption that the three-phase respiratory pattern is generated through interactions among a small number of functional populations, each characterized by its discharge pattern and excitatory or inhibitory output. For the present reconstruction, we assume that the phase-dependent synaptic inputs originate within the same five respiratory populations from which the receiving cells were sampled. Too few recorded neurons were identified by transmitter to compare inputs to excitatory and inhibitory cells within a discharge pattern. We therefore grouped receiving cells by discharge pattern alone. The number and anatomical subdivisions of the underlying functional populations remain open biological questions.

Within this description, synaptic inputs to a recorded neuron provide a postsynaptic readout of convergent activity from functional populations. A recorded cell can reveal excitation received while it is silent, or inhibition received while it is active, neither of which can be inferred from its discharge alone. Separating excitatory and inhibitory conductances has constrained circuit mechanisms in sensory and rhythmic motor systems (Wehr and Zador, 2003; Atallah and Scanziani, 2009; Berg et al., 2007; Guzulaitis and Hounsgaard, 2017; Berg, 2018).

Previous approaches to inferring functional interactions in active respiratory circuits include simultaneous multisite extracellular recordings and correlation analysis of neuronal spike trains (Segers et al., 2008). These approaches have informed models of respiratory circuitry (Balis et al., 1994; Lindsey et al., 2012). Paired extracellular and intracellular recordings have also been used to infer synaptic connections by averaging intracellular membrane potentials triggered by extracellular spikes to reveal postsynaptic events (Tian et al., 1999; Duffin et al., 2000). Our previously developed conductance-inference method reconstructs excitatory and inhibitory conductance profiles throughout the respiratory cycle from single sharp-microelectrode recordings of neurons with different discharge patterns in the pre-BötC and BötC of mature rat preparations in situ (Molkov et al., 2025). Appendix Figure A1 illustrates the progression from intracellular recordings to conductance profiles. In a separate study, Molkov et al. (2026a) extended conductance analysis to transmitter-identified neurons in rhythmic pre-BötC slices.

The conductance profiles from Molkov et al. (2025) suggested functional circuit connectivity motifs that could be combined into a functional connectome of the active respiratory network. These included aug-E excitation of pre-I/I and inspiratory excitation of expiratory neurons. Such assignments relate received synaptic input to presynaptic activity, assuming that a presynaptic population’s contribution follows its discharge pattern. Each conductance, however, combines inputs from many neurons, and candidate presynaptic populations are active during overlapping portions of the respiratory cycle. Different combinations of active populations can therefore produce similar input time courses. Distinguishing their contributions requires comparing these alternatives while accounting for variation among cells in the receiving population.

Here we analyze many of the same intracellular recordings together with additional recordings from the archive. We use this larger sample to quantify input contributions and distinguish their possible presynaptic origins. We assume that neuronal population discharge patterns reported independently by Smith et al. (2007) represent presynaptic timing in the recorded preparation.

We also assume that each sample of receiving cells represents the corresponding discharge-pattern population. We have developed statistical tests to determine which candidate inputs are supported in its mean conductance and which presynaptic assignments remain unresolved (Section 2.7; Appendix A.3). Combining the supported inputs to each receiving population gives a directed candidate functional connectome. Finally, we compare this circuit with the three-phase architecture developed to explain rhythmic respiratory pattern generation and its transformations after brainstem transections, identifying both shared interactions and additional connections supported by the synaptic inputs.

## 2 Methods

### 2.1 Ethical approval

We reanalyzed intracellular recordings from the same archive used by Molkov et al. (2025); no new animal experiments were performed. The Animal Care and Use Committee of the National Institute of Neurological Disorders and Stroke approved the original procedures (Animal Study Proposal 1154–21). The NIH intramural animal care program follows the U.S. Public Health Service Policy on Humane Care and Use of Laboratory Animals.

### 2.2 From recorded cells to population connections: approach overview

Figure 1 outlines how we infer circuit connections from cellular conductances and independently recorded population activity. We grouped recorded cells by respiratory discharge pattern and assumed that each sample represents the corresponding receiving population. Candidate presynaptic populations are defined jointly by discharge pattern and excitatory or inhibitory output. We assume that the phase-dependent inputs originate within the five represented discharge-pattern populations and that their activity templates describe presynaptic timing.

**Figure 1.**
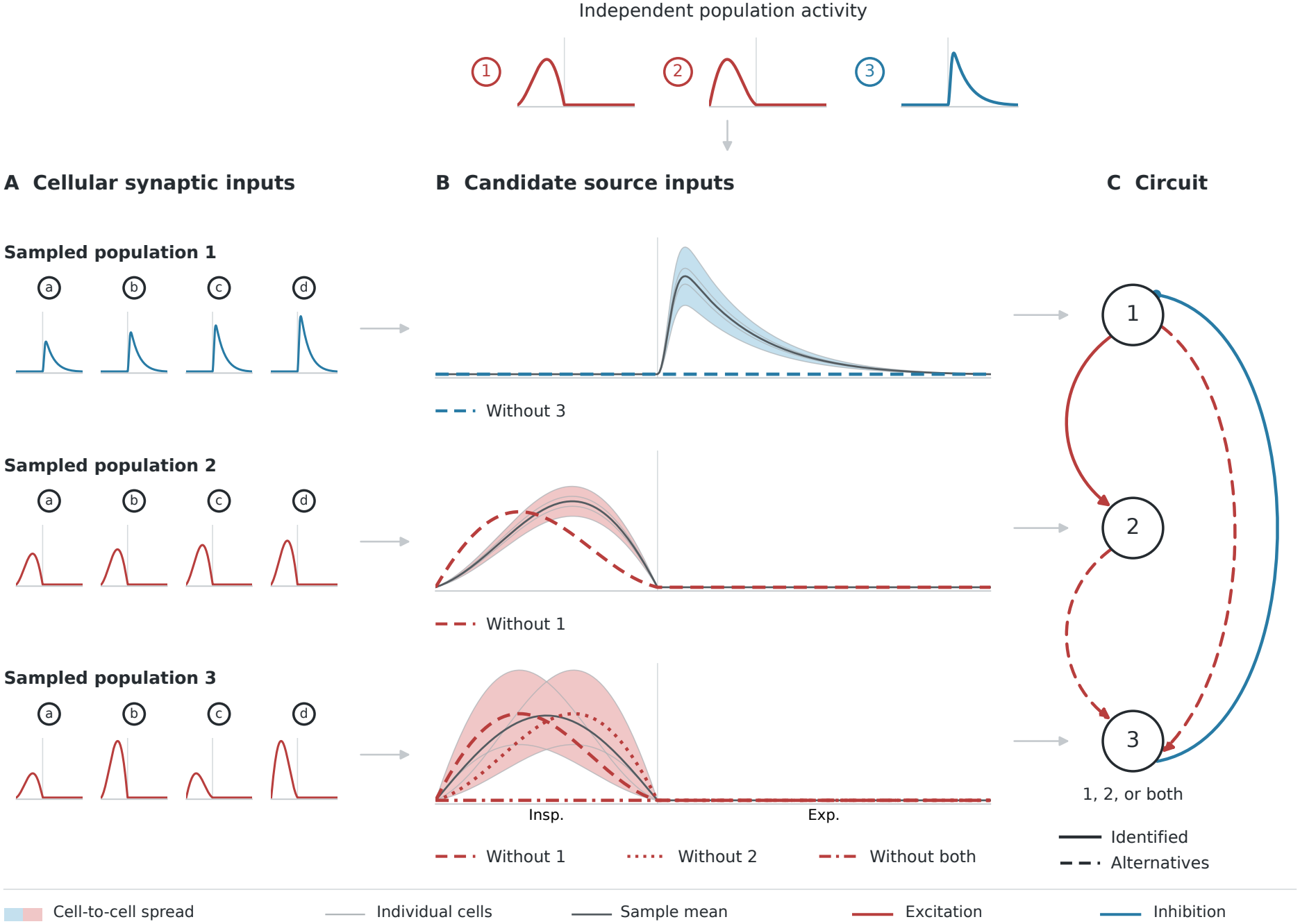
From individual cells to a population circuit. A, cells a–d are grouped by discharge pattern. Profiles show inhibitory conductance in population 1 and excitatory conductance in populations 2 and 3 over the respiratory cycle. B, candidate source activity patterns appear above. Grey curves repeat the cellular profiles; charcoal curves show sample means and shading shows cell-to-cell ranges. Broken curves show fits omitting the indicated sources. The bottom row allows excitation from population 1, population 2, or both; omitting both leaves no excitatory input. C, solid connections indicate individually distinguished inputs; dashed connections indicate unresolved alternatives. Red arrowheads denote excitation and filled blue circles inhibition. Vertical guides mark inspiratory offset. All panels are schematic. Shading describes variation; statistical support comes from across-cell tests (Section 2.7).

We approximated each reconstructed conductance as a nonnegative sum of contributions proportional to the candidate source activities. Excitatory and inhibitory conductances were fitted separately. Similar source timing can permit several explanations of the same input profile. We therefore tested whether the population mean conductance could be explained after omitting a source, using variation among cells to estimate uncertainty. Rejection supports that source’s contribution. Rejection of a joint omission without rejection of either individual omission supports at least one member, while allowing both to contribute.

The supported inputs define a candidate circuit between discharge-pattern populations (Figure 1C). Unresolved sources remain alternatives. Section 2.7 describes the tests, and Appendix A.3 gives their derivation and a worked example.

### 2.3 Experimental preparation and recordings

Experiments used in situ arterially perfused brainstem-spinal cord preparations from male Sprague-Dawley rats, 3–4 weeks old and weighing 45–70 g. Rats were deeply anesthetized with 5% isoflurane and decerebrated. The preparation was maintained at 31^*°*^C in carbogenated Ringer solution, with neuromuscular transmission blocked by vecuronium. Full preparation and histological procedures are described by Molkov et al. (2025).

Phrenic and central vagal motor output were recorded with suction electrodes. Sharp intracellular recordings were obtained from neurons in the pre-BötC and BötC using 45–70 MΩ electrodes and a dual current-clamp/voltage-clamp amplifier. Injected current or holding voltage was varied across respiratory cycles to obtain phase-specific current-voltage relations. Hyperpolarization reduced contamination by voltage-dependent currents.

### 2.4 Recording selection and sampled populations

We analyzed 168 recordings from 150 cells (Molkov et al., 2026b): 17 pre-I/I, 41 ramp-I, eight early-I, 54 post-I, and 30 aug-E cells. There were 150 current-clamp and 18 voltage-clamp recordings. Recordings required stable intracellular measurements, sufficient variation in injected current or holding voltage, and identifiable inspiratory onset and offset. We selected the longest eligible contiguous interval with a respiratory-period coefficient of variation no greater than 15%, before fitting candidate inputs. This broadens the earlier criteria of variation below 10% and at least three current levels maintained for five cycles each (Molkov et al., 2025).

Discharge-pattern assignments followed archive annotations and review of respiratory timing and membrane-potential modulation. Fifty-one recordings from 46 cells retained provisional assignments because downsampling or hyperpolarization limited verification. Cells from the pre-BötC and BötC regions were grouped by discharge pattern rather than location. The archive does not document blinded classification.

Repeated recordings were averaged within clamp mode and then across modes to give each cell equal weight. The available records do not link each cell to its animal or preparation. We treated cells as independent sampling units; dependence among cells from a shared preparation remains unquantified. Cells recorded on the same day may have come from the same preparation (see Section 2.8).

### 2.5 Phase alignment and conductance reconstruction

Phrenic activity provided the respiratory reference in all recordings. Inspiratory onset and offset were defined by 20% rising and 50% falling crossings of burst amplitude. We rescaled inspiration and expiration separately onto a common cycle of 100 phase bins, placing inspiratory offset at approximately phase 0.24. This common phase coordinate permits comparison between recordings with different cycle durations.

After median filtering to attenuate spikes, we used the current-voltage relation at each respiratory phase to reconstruct the time courses of excitatory and inhibitory conductance (Molkov et al., 2025). Appendix A.1 gives the equations and an illustrated example.

The dynamic excitatory and inhibitory conductances, *g*_E_ and *g*_I_, are each reconstructed conductance’s variation above its cycle minimum. Tonic synaptic conductance and leak remain combined. We divided both profiles from each recording by the larger of their two peak values, preserving the relative amplitudes of excitation and inhibition. Cross-cell comparisons use this normalized scale because uniform physical-unit calibration is not established across the archive.

### 2.6 Candidate presynaptic activity and individual fits

We digitized five activity profiles from the pontine-medullary condition in Figure 4 of Smith et al. (2007): pre-I/I, ramp-I, early-I, post-I, and aug-E. Each activity template was aligned to the respiratory reference, minimum-subtracted, and peak-normalized. We aligned the post-I template’s rising half-amplitude point to the published inspiratory-offset guide, assuming that its apparent displacement arose in the original figure. The same template was available for excitatory and inhibitory sources, assuming common timing for either output type within a discharge-pattern population.

For each recording and conductance type, we fitted the profile as a nonnegative weighted sum of the five activity templates. We chose the weights to minimize the summed squared fitting error over the full cycle (non-negative least squares, NNLS; Lawson and Hanson, 1995), without an offset. Appendix A.2 gives the normalization and fitting equations. This additive representation is an assumption about the processed, minimum-subtracted waveforms. Each fitted contribution’s area was divided by the total area of fitted excitation and inhibition in that recording. Fractions were averaged within clamp mode, within cell, and equally across cells. Figures show individual cell profiles and averages of individual fits.

**Table 1:** Inspiratory input combinations consistent with the omission tests.

| Target | Excitatory input | Inhibitory input |
|---|---|---|
| pre-I/I | (pre-I/I + early-I) or (ramp-I + early-I) | pre-I/I or ramp-I or early-I |
| ramp-I | (pre-I/I + ramp-I) or (ramp-I + early-I) | early-I |
| early-I | pre-I/I or (ramp-I + early-I) | No inspiratory input established |
| post-I | pre-I/I or ramp-I | (pre-I/I + early-I) or (ramp-I + early-I) |
| aug-E | pre-I/I or early-I | (pre-I/I + early-I) or (ramp-I + early-I) |

A plus sign joins inputs retained together; “or” separates alternatives. Removing any member from a listed combination conflicts with at least one rejected omission. Additional inputs remain possible. Both expiratory templates remain available.

### 2.7 Statistical evidence and inspiratory alternatives

We tested whether a population mean conductance could be explained without specified candidate inputs, allowing all retained templates to contribute with nonnegative weights. For each omitted template, we constructed a timing contrast from the residual after approximating it with the retained templates. These weights depend only on the activity templates. Applying the same contrast to every cell gives one score per cell. Under the omission hypothesis, the expected score is nonpositive; a sufficiently positive sample mean supports rejection. We compared the mean score with zero using a one-sided *t* test and the across-cell standard error. Appendix A.3 derives the procedure.

For each receiving population and conductance type, we tested all seven nonempty omissions of the three inspiratory templates, with both expiratory templates retained. Post-I and aug-E omissions were tested individually. We combined Bonferroni correction within each omission, closed testing across the seven inspiratory omissions (Marcus et al., 1976), and correction across ten inspiratory families and twenty expiratory tests. An adjusted P value below 0.05 supports the omitted input or set. The 30 overview comparisons comprise whole-inspiratory, post-I, and aug-E omissions for each population and conductance type.

Rejection of an individual omission distinguishes that input. Rejection of a joint omission establishes that at least one member contributes. For unresolved inspiratory inputs, we report retained combinations from which no member can be removed while remaining consistent with all rejected omissions. Combinations containing them also remain possible. Failure to reject leaves the omitted input’s presence unresolved.

The tests assume independent, representative cells, fixed activity templates, and a population mean expressible as a nonnegative sum of those templates. Exact *t*-test significance levels require normally distributed cell scores with common positive variance. Phase bins within a cell may be correlated. Several score distributions were skewed, making significance levels approximate, especially for the eight-cell early-I sample. Across-cell uncertainty describes cellular variation conditional on the templates and reconstruction.

### 2.8 Sensitivity to phase alignment, individual cells and recording days

We assessed timing sensitivity by aligning activity templates to each recording’s 40% or 60% falling phrenic landmark in place of the 50% landmark. To assess the influence of individual cells, we repeated the omission tests after removing one cell at a time. We identified recording dates from the file names. To give each recording day equal weight within a discharge-pattern population, we averaged the cell scores for each day and repeated the 30 grouped omission tests using those daily means. Processing settings, the recording audit, and complete statistical and diagnostic outputs accompany the data and code.

## 3 Results

### 3.1 Synaptic inputs vary within and between discharge-pattern populations

Reconstructing the circuit requires both the activity of candidate presynaptic populations and the synaptic inputs received by the recorded neurons. Figure 2A shows the five activity templates from Smith et al. (2007). pre-I/I, ramp-I, and early-I overlap during inspiration but differ near its onset and termination. The declining post-I and increasing aug-E profiles overlap during expiration.

**Figure 2.**
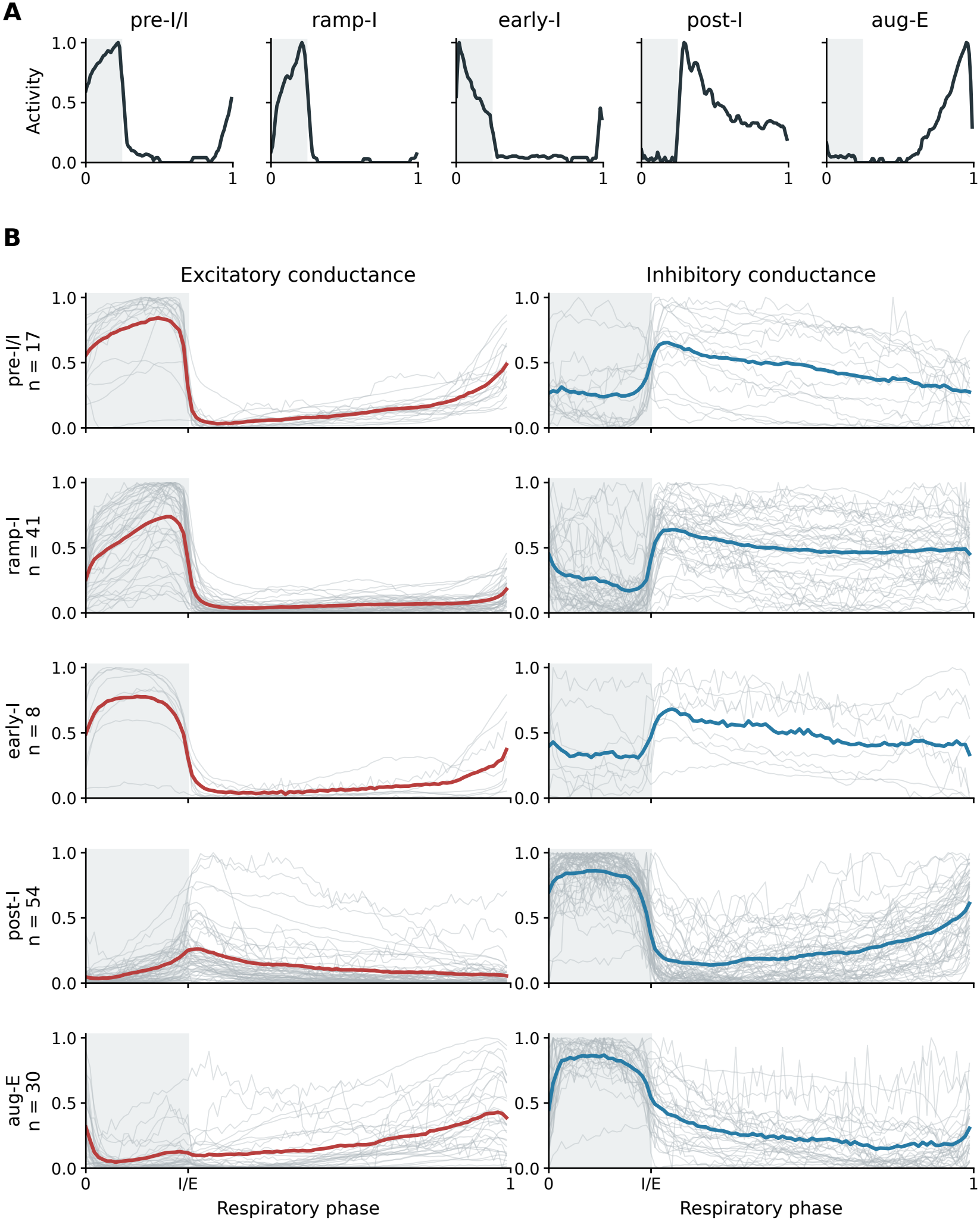
Candidate discharge patterns and reconstructed synaptic inputs. A, activity templates digitized from the pontine-medullary preparation in Smith et al. (2007), aligned to a common cycle, minimum-subtracted, and peak-normalized. B, dynamic excitatory and inhibitory conductances reconstructed in the present analysis. Grey curves represent individual cells; colored curves are sample means. Both conductances share the larger of their two peak values in each recording as normalization. Samples contain 17 pre-I/I, 41 ramp-I, eight early-I, 54 post-I, and 30 aug-E cells. Grey shading marks inspiration; I/E marks the inspiratory-expiratory transition.

We reconstructed conductance profiles from 150 cells (Figure 2B). For each recording, the excitatory and inhibitory profiles share a scale, preserving their relative amplitudes. Cells within each discharge-pattern population varied substantially in the magnitude and duration of their inputs.

Inspiratory populations showed prominent excitation during inspiration and inhibition extending into expiration. In the sample means, excitatory conductance increased during inspiration in ramp-I neurons and during late expiration in aug-E neurons. Increasing excitation therefore accompanied the active phase of both discharge patterns. Expiratory populations also received excitation during inspiration, when they were silent, alongside stronger inhibition. We used the five activity templates to separate these conductances into candidate presynaptic contributions, fitting each recording individually.

### 3.2 Candidate excitatory and inhibitory contributions to the recorded conductances

Figure 3 shows the mean individual fits and their combined inspiratory, post-I, and aug-E contributions. The inspiratory curve sums the separately fitted pre-I/I, ramp-I, and early-I inputs. Contribution magnitudes are expressed as percentages of the total area under the fitted excitatory and inhibitory conductance curves.

**Figure 3.**
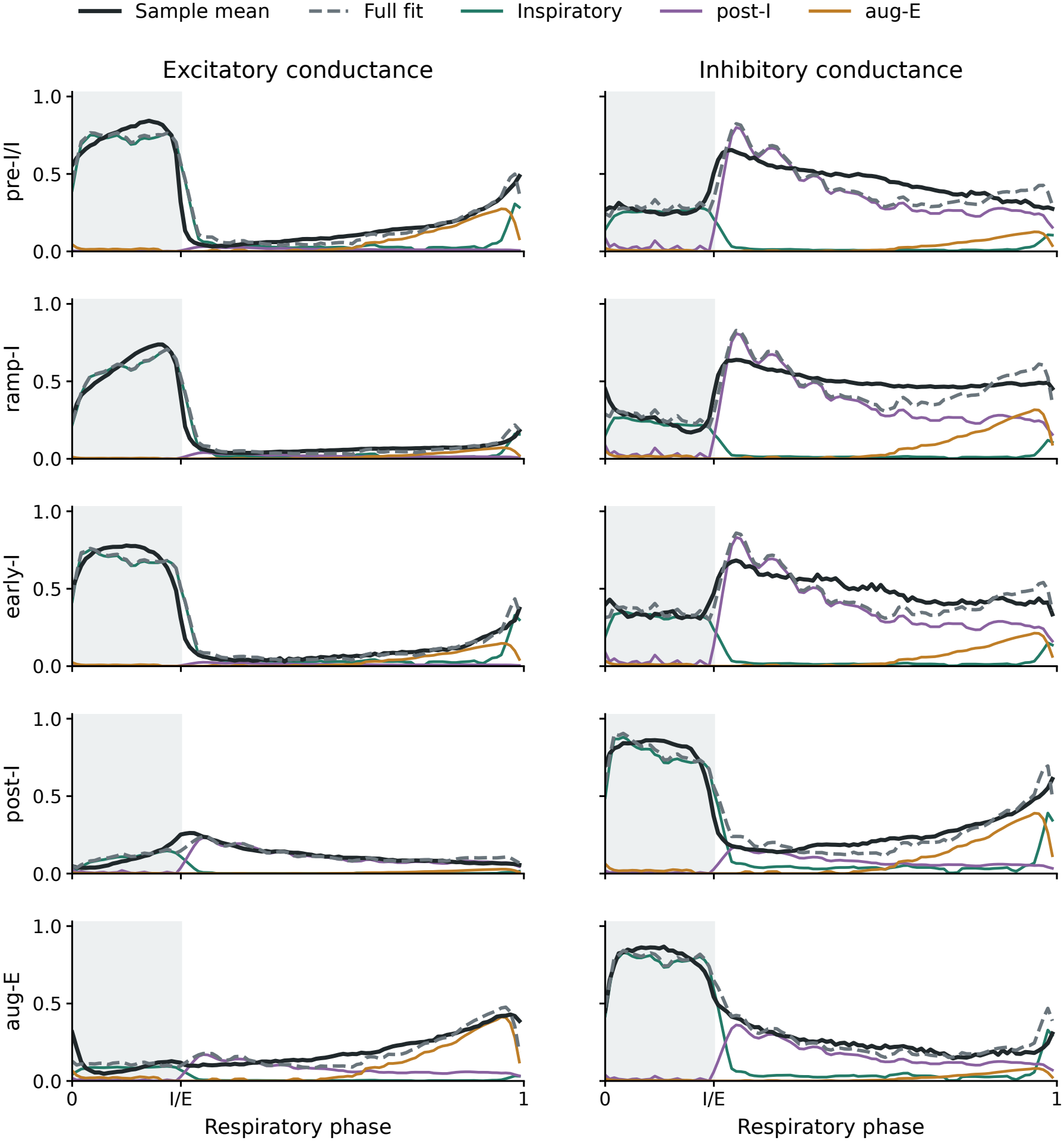
Conductance fits and candidate presynaptic contributions. Black curves show sample mean conductances; dashed grey curves show mean individual NNLS fits. Green, purple, and ochre curves show combined inspiratory, post-I, and aug-E contributions. All five templates are fitted jointly to each conductance, with the three inspiratory contributions summed for display. All fitted components are shown, including those without statistical support. Cells receive equal weight; normalization follows Figure 2. Grey shading marks inspiration.

In pre-I/I and ramp-I, post-I activity matches the declining expiratory inhibition and contributes 40.2% and 44.4% of the total fitted conductance area, respectively. An additional aug-E component contributes inhibition later in expiration in ramp-I, with a fitted share of 10.2%.

The late-expiratory excitation of pre-I/I includes an aug-E contribution, with a mean fitted share of 8.9%. The post-I and aug-E populations receive smaller inspiratory excitatory contributions, 5.5% and 4.3%, respectively, alongside their prominent inspiratory inhibition.

The five-template fits approximate the principal temporal features. Median cell-level root-mean-square errors range from 3.2% to 8.8% of the normalization peak for excitation and from 9.8% to 13.2% for inhibition across populations. Supporting Data S1 gives the number of cells and the distribution of fitted contributions for each population, together with the complete omission-test results.

### 3.3 Statistical evidence constrains candidate circuit connections

The fitted contributions describe possible origins of the conductances in Figure 3. To determine which inputs could be distinguished, we tested explanations that omitted each candidate contribution while allowing the other activity patterns to contribute (Section 2.7). We found statistical support for 20 of the 30 combined-inspiratory, post-I, and aug-E inputs: nine excitatory and eleven inhibitory. For these inputs, we rejected the hypothesis that the remaining patterns could explain the population mean conductance.

We assembled the supported incoming connections into the directed functional circuit in Figure 4A, under the source and sampling assumptions illustrated in Figure 1C. Each connection starts at a candidate excitatory or inhibitory presynaptic population and ends at a broader receiving discharge-pattern population. Excitatory and inhibitory candidate sources are shown separately within each enclosure. Incoming connections terminate on the enclosure and describe the mean input to cells with that discharge pattern.

**Figure 4.**
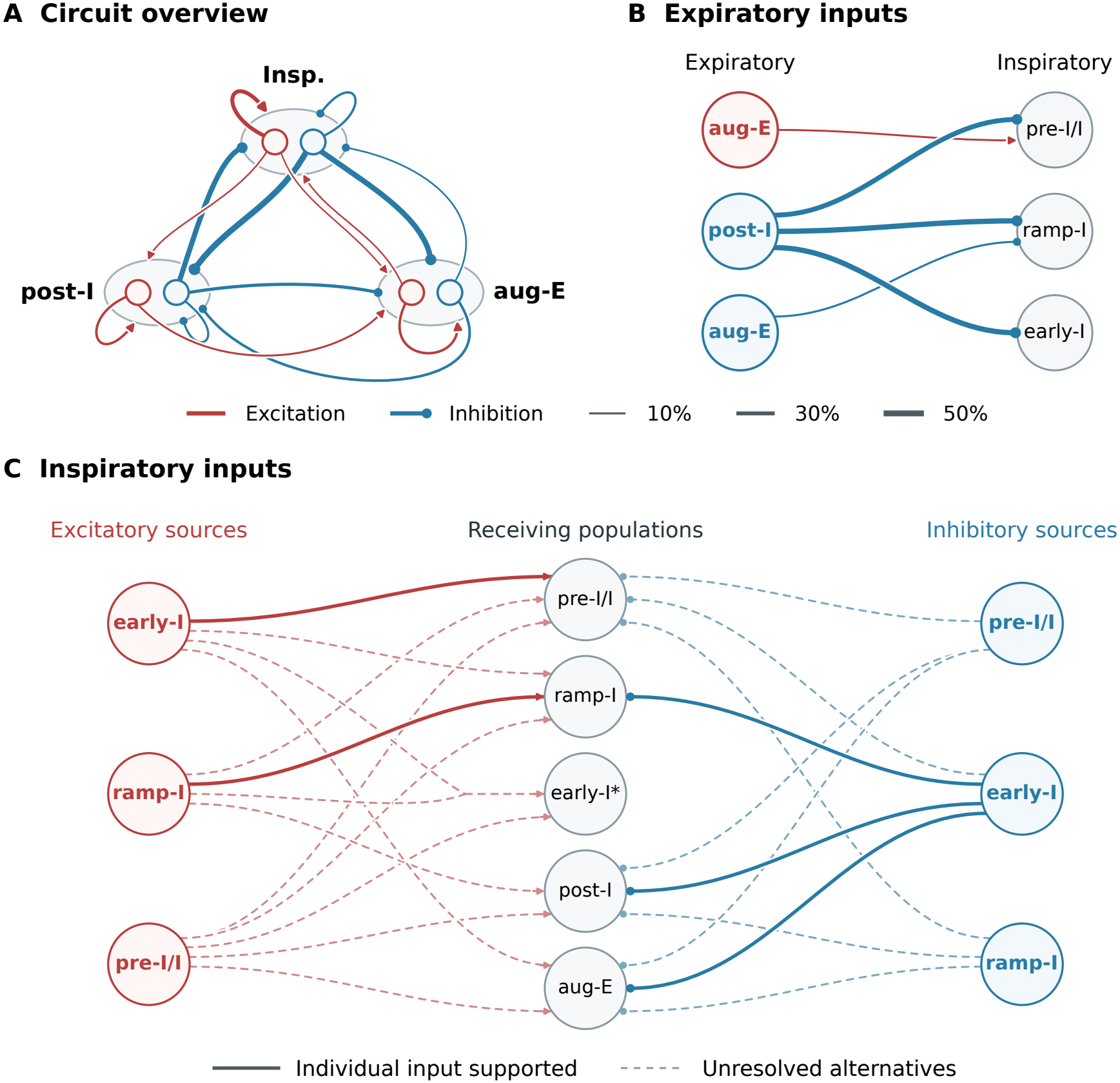
Conductance-constrained candidate circuit. Red arrowheads denote excitation and filled blue circles inhibition. Paired nodes represent candidate excitatory and inhibitory sources; enclosing targets pool cells by discharge pattern. A, overview combining pre-I/I, ramp-I, and early-I as Insp.; an incoming connection has support in at least one constituent population. B, expiratory inputs to individual inspiratory populations. In A and B, widths show mean fitted percentages of total dynamic excitatory plus inhibitory conductance area. For the combined inspiratory target, widths average the three population means equally, including unsupported fitted contributions. C, individually distinguished inputs (solid) and unresolved alternatives (dashed; Table 1). For each target, at least one dashed input of each displayed color is needed alongside any solid input. *Early-I excitation: pre-I/I alone or ramp-I and early-I together (merged branches). Connections summarize separate omission tests.

The overview combines the three inspiratory populations in one enclosure. An incoming connection to this enclosure indicates support in at least one constituent population. Figure 4B identifies the specific inspiratory populations receiving expiratory inputs. Post-I inhibition is supported in pre-I/I, ramp-I, and early-I. Aug-E inhibition is additionally supported in ramp-I, and aug-E excitation is supported in pre-I/I.

Both expiratory populations receive inhibition from inspiratory populations and from the other expiratory population. Post-I-pattern inhibition of post-I cells is also supported. Excitation with post-I timing is supported in both post-I and aug-E populations; excitation with aug-E timing is supported in aug-E. Connections returning to a population denote candidate interactions among neurons with the same discharge pattern.

### 3.4 Some inspiratory inputs can be assigned to individual discharge patterns

Differences in timing among the overlapping inspiratory patterns may distinguish individual contributions. Where we cannot distinguish them, we retain alternative presynaptic origins of a supported input. By testing all inspiratory subsets, we distinguished five individual discharge-pattern contributions across the ten population/conductance combinations. Figure 4C shows these individual inputs and the remaining alternatives. Table 1 lists the combinations consistent with the tests. Each is minimal in the sense that removing another member violates at least one rejected omission; larger combinations remain possible.

For ramp-I excitation, we rejected omission of the ramp-I pattern, but also excluded ramp-I alone as the inspiratory contribution. The remaining combinations included either pre-I/I or early-I in addition to ramp-I. For pre-I/I excitation, we found support for an early-I-pattern contribution together with either pre-I/I or ramp-I. These assignments depend on temporal features within the chosen activity patterns.

For early-I excitation, the retained alternatives were pre-I/I alone or ramp-I and early-I together. Thus, inspiratory excitation was supported without identifying any one of the three patterns individually. The converging dashed branches in Figure 4C denote the two-pattern alternative, not two separately established connections.

Post-I excitation illustrates the distinction between a dominant fitted contribution and a distinguishable presynaptic input. Ramp-I provided the largest fitted inspiratory excitatory contribution in 50 of 54 cells. We found statistical support for inspiratory excitation but could not assign it uniquely to pre-I/I or ramp-I. Figure A2 in the Appendix shows these alternative explanations and the resulting connections. The dashed inputs to post-I in Figure 4C therefore retain both candidate origins, including their possible co-occurrence.

Early-I-pattern inhibition was individually supported for ramp-I, post-I, and aug-E. In the two expiratory populations, the retained combinations also included a pre-I/I or ramp-I inhibitory contribution. No inspiratory inhibitory input was statistically established for the eight-cell early-I sample.

Changing the activity-template inspiratory offset to either nearby phrenic landmark (Section 2.8) left all omission decisions unchanged.

Nineteen of the 20 supported grouped inputs retained support after every single-cell removal. Inspiratory inhibition of pre-I/I was the exception: its adjusted P value ranged from 0.020 to 0.055 and crossed 0.05 after four of 17 removals. All five individually distinguished inspiratory contributions and 29 of the 32 rejected inspiratory-subset omissions retained support after every removal.

The 150 cells were recorded on 64 days. After averaging within recording day, 19 of the 20 supported grouped inputs retained support; the exception was again inspiratory inhibition of pre-I/I (adjusted *P* = 0.112).

### 3.5 Additional excitation beyond the transection-based circuit

Smith et al. (2007) mapped respiratory discharge patterns along the ventral respiratory column. Drawing on these recordings and earlier physiological and modeling studies (Rybak et al., 1997; 2004; 2007), they proposed a three-phase circuit to explain how the respiratory pattern changed after sequential brainstem transections. Figure 5 summarizes its connections after grouping neurons by discharge pattern, allowing comparison with the conductance-based circuit in Figure 4.

**Figure 5.**
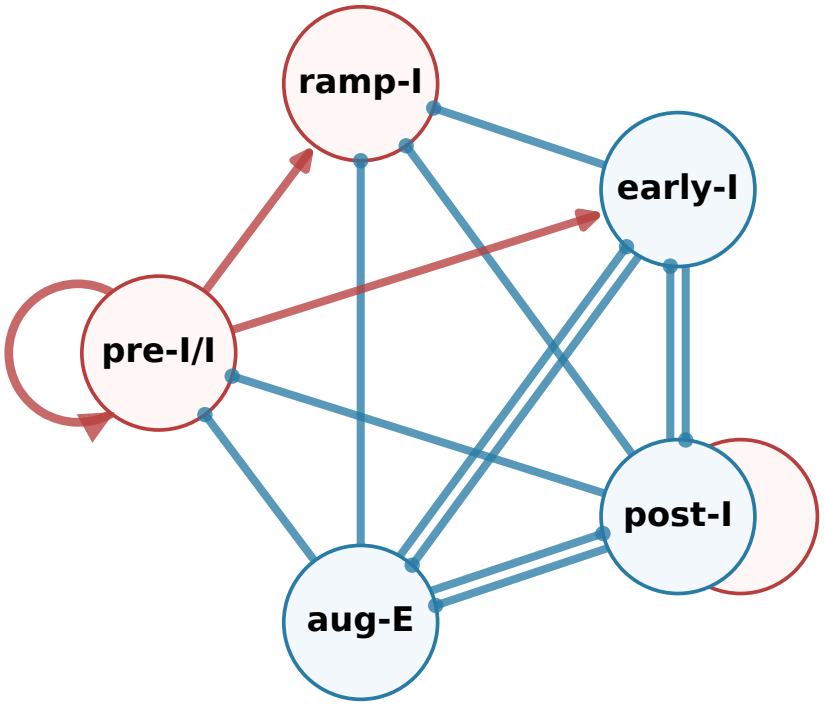
Transection-based three-phase circuit. Connections from Figure 7 and Table 3 of Smith et al. (2007), grouped by discharge pattern. Red arrowheads denote excitation and filled blue circles inhibition. Early-I combines early-I(1) and early-I(2); the post-I pair includes excitatory post-I(e) premotor neurons and shares the displayed incoming connections.

Twelve of the fourteen connections in Figure 5 have support at the combined-inspiratory resolution of Figure 4. Nine inhibitory connections are individually supported. These include post-I inhibition of all three inspiratory populations, early-I inhibition of ramp-I and both expiratory populations, and reciprocal post-I/aug-E inhibition. The three excitatory projections from pre-I/I match supported inspiratory inputs, although their assignment specifically to pre-I/I is unresolved. Aug-E inhibition of pre-I/I and early-I, present in the earlier circuit, is not statistically established here.

Relative to Figure 5, the conductance-based circuit contains six additional excitatory inputs: aug-E excitation of pre-I/I; inspiratory and post-I excitation of post-I; and inspiratory, post-I, and aug-E excitation of aug-E. It also contains two additional inhibitory inputs, from inspiratory populations to pre-I/I and from post-I to post-I. The additional excitation therefore includes inputs during both the active and silent phases of expiratory neurons. Several of these motifs were proposed from conductance timing in Molkov et al. (2025); here their fitted magnitudes and omission-test evidence are quantified across the expanded sample.

For this comparison, we grouped neurons more broadly than in the original anatomical circuit. Its ramp-I population represents premotor neurons in the rostral ventral respiratory group (rVRG), whereas we grouped ramp-I recordings by timing rather than location. We also combined its two early-I populations in Figure 5. At finer temporal resolution, we rejected pre-I/I as the sole inspiratory excitatory input to pre-I/I and ramp-I. Thus, the earlier circuit agrees with major inhibitory features but needs additional inputs to represent the reconstructed conductances using these activity templates.

## 4 Discussion

Our reconstructed synaptic inputs and novel functional circuit architecture of the respiratory network retain the major inhibitory interactions of the previous three-phase respiratory activity circuit and add phase-dependent excitation not present in that circuit. In three-phase activity circuit concepts, inhibitory synaptic interactions among early-I, post-I, and aug-E neurons switch the network between inspiration, post-inspiration, and late expiration (Richter, 1982; Rybak et al., 1997; 2004; 2007; Smith et al., 2007). We have found support for the inhibitory inputs on which each of these transitions depends, and post-I-pattern inhibition provides the largest fitted contribution to the conductances of all inspiratory populations. In the circuit of Smith et al. (2007), phase-dependent excitation originates from pre-I/I neurons and reaches only inspiratory populations, whereas expiratory populations receive tonic excitatory drive. This original depiction of circuit interactions was based on limited information on actual synaptic interactions. Our analysis of the archived recordings supports excitation with other timing and other targets.

Excitatory input increased during ramp-I and aug-E discharge and occurred in pre-I/I neurons during late expiration and in both expiratory populations during inspiration, when they are silent. Our previous conductance study suggested these excitatory inputs (Molkov et al., 2025); here we tested each against alternative presynaptic timing across cells in the expanded sample. We propose that phase-dependent excitation acts together with inhibition in shaping ramp-I and aug-E discharge and initiating inspiration.

### 4.1 Inhibitory inputs and respiratory phase transitions

We have found support for post-I inhibition of all three inspiratory populations, early-I inhibition of ramp-I and both expiratory populations, aug-E inhibition of ramp-I, and inhibition in both directions between post-I and aug-E (Figure 4). Several of these inputs correspond to synaptic potentials recorded intracellularly in cats. Ballantyne and Richter (1984) described declining early-inspiratory and augmenting expiratory inhibitory potentials in bulbar inspiratory neurons. Ballantyne and Richter (1986) described inspiratory phase inhibition of expiratory bulbospinal neurons and, in some of them, decrementing post-inspiratory inhibition. In cats and rats, spike-triggered averaging showed that BötC neurons with augmenting expiratory discharge monosynaptically inhibit inspiratory bulbospinal neurons (Jiang and Lipski, 1990; Tian et al., 1999). Our reconstruction quantifies these inhibitory patterns over the full cycle in five discharge-pattern populations and tests each against alternative presynaptic activity timing.

Three-phase activity circuit concepts mechanistically assign these inputs to specific phase transitions. Richter (1982) proposed that post-inspiratory inhibition gates the inspiratory neuron network immediately after inspiration ends. In the model of Smith et al. (2007), early-I, post-I, and aug-E populations form a ring of mutual inhibition. Declining early-I inhibitory neuron activity during inspiration releases post-I inhibitory neurons, which terminate inspiration.

Declining post-I activity then releases aug-E neurons. Before the end of expiration, the excitatory pre-I/I population is released from expiratory inhibition and initiates the next inspiration. Rubin et al. (2009b) showed how adaptation, release from inhibition, and recruitment of suppressed populations generate these transitions in the model. The inputs supported here include those on which each transition depends: declining early-I inhibition of post-I for inspiratory termination, post-I inhibition of aug-E for the transition to late expiration, and post-I inhibition of pre-I/I, which delays inspiratory onset.

Regional perturbations show that inhibition in the pre-BötC and BötC can alter respiratory timing. Glycinergic pre-BötC neuron stimulation terminated inspiration or delayed its next onset according to stimulus phase (Sherman et al., 2015). In optogenetic experiments in transgenic mouse in situ preparations, Ausborn et al. (2018) found that inhibitory pre-BötC neuron photostimulation accelerated or slowed the respiratory rhythm depending on intensity, whereas BötC inhibitory neuron photostimulation slowed or suppressed the rhythm. Local pharmacological blockade of fast synaptic inhibition did not disrupt rhythmic breathing in the adult rats studied by Janczewski et al. (2013), whereas Marchenko et al. (2016) found regional effects of pharmacological disruption of synaptic inhibition on phase duration and pattern, including loss of rhythmic output in some preparations. In our recordings, post-I-pattern inhibition was the largest fitted contribution to the conductances of each inspiratory population (Figure 3). In the model of Rubin and Smith (2019), strengthening this input prolonged expiration.

### 4.2 Late-expiratory inputs and inspiratory onset

In the circuit of Smith et al. (2007), aug-E neurons inhibit pre-I/I, early-I, and ramp-I neurons during late expiration (Figure 5). We found support for aug-E-pattern inhibition of ramp-I, matching the augmenting expiratory inhibition recorded in bulbar inspiratory neurons (Ballantyne and Richter, 1984). Aug-E-pattern inhibition of pre-I/I and early-I was not established from our conductance reconstructions. For pre-I/I, the supported aug-E-pattern input was excitatory (Figure 4B). The reconstructed pre-I/I inputs therefore combine declining post-I inhibition with excitation that rises through late expiration (Figure 3).

We hypothesize that these two inputs cooperate in recruiting the next inspiratory phase. In the model of Smith et al. (2007), the pre-I/I population, which is critical for inspiratory rhythm generation, is released from expiratory inhibition before ramp-I neurons and initiates inspiration. Increased excitatory drive to this population advanced inspiratory onset in the model of Rubin and Smith (2019). Excitation that rises during late expiration would act as such a drive at the phase when the next inspiration is recruited. Inspiratory onset would then depend on a late-expiratory excitatory input in addition to release from inhibition. We did not test whether late-expiratory inputs differ between pre-I/I and ramp-I neurons.

The pre-I/I discharge pattern is that of the excitatory pre-BötC population proposed as the kernel population for inspiratory rhythm generation. The pre-I/I neurons reported by Molkov et al. (2025) were recorded in the pre-BötC; recording sites are not documented for all cells in the expanded sample, but for pre-I/I cells with documented locations, all were in the pre-BötC. This population and other inspiratory neurons remain rhythmic after isolation from more rostral compartments (Smith et al., 2007; Rubin et al., 2009b). Rubin and Smith (2019) obtained similar simulated three-phase rhythms when this population was intrinsically bursting or tonically active in isolation. In a separate conductance reconstruction study of transmitter-identified pre-BötC neurons in rhythmic slices, Molkov et al. (2026a) found a self-exciting excitatory inspiratory population, including pre-I/I neurons, coupled to inspiratory and expiratory inhibitory populations. Inspiratory neurons in those slices received tonic inhibition during expiration. In the present in situ recordings, the dynamic expiratory inputs to pre-I/I neurons followed distinct post-I and aug-E patterns. The in situ recordings are consistent with recurrent excitation of pre-I/I neurons, although we could not distinguish a pre-I/I contribution from a ramp-I contribution to their inspiratory excitation (Table 1). The in situ vs. in vitro preparations differ in species, age, excitability, and retained inputs. Burst termination and recovery within an excitatory inspiratory network have been attributed to activity-dependent processes, including synaptic depression (Rubin et al., 2009a; Kottick and Del Negro, 2015). If these processes operate in the three-phase rhythm, they do so alongside expiratory inputs that first inhibit and then excite pre-I/I neurons.

### 4.3 Excitation and the formation of discharge patterns

Sample mean excitatory conductance increased during ramp-I and aug-E discharge (Figure 2B), with supported contributions matching each population’s augmenting discharge. Two mechanisms have been proposed for the inspiratory ramp. Richter (1982) proposed a ramp-generating loop of mutually exciting inspiratory neurons. In the models of Rybak et al. (2007) and Smith et al. (2007), ramp-I neurons receive excitation from pre-I/I neurons, and declining early-I inhibition shapes their augmenting discharge. We found support for the synaptic input required by each mechanism: early-I-pattern inhibition of ramp-I, and ramp-I-pattern excitation that the other inspiratory patterns could not replace. The synaptic input to ramp-I neurons therefore combines growing excitation with declining inhibition. Their relative contributions to the discharge ramp remain to be determined.

Studies of the pre-BötC separate the timing of inspiratory events from the formation of the inspiratory burst (Kam et al., 2013; Cui et al., 2016), and Ashhad and Feldman (2020) showed how increasingly synchronous convergent excitation recruits inspiratory output neurons. Inhibition during inspiration also shapes the burst: limiting inspiratory depolarization shortened the subsequent refractory period (Baertsch et al., 2018), and inhibitory subpopulations differed in their effects on burst onset versus duration and amplitude (Chang et al., 2024). The rising excitation and early-I-pattern inhibition of ramp-I neurons that we identified are inputs through which such control could act during the three-phase rhythm.

The identified inputs to aug-E neurons likewise combine declining inhibition with rising excitation. In the model of Smith et al. (2007), expiratory populations receive tonic excitatory drive, and aug-E discharge begins as post-I inhibition declines. We found support for post-I inhibition of aug-E and for excitation with aug-E timing. Jiang and Lipski (1990) identified excitatory potentials in two expiratory bulbospinal neurons by spike-triggered averaging from BötC augmenting neurons, alongside widespread inhibitory connections. Within our five discharge-pattern representation, the supported inputs to aug-E and pre-I/I neurons indicate excitatory neurons with aug-E discharge, a novel feature of our reconstructed circuit relative to that of Smith et al. (2007).

We found support for inspiratory excitation of post-I and aug-E populations during a phase in which these neurons are normally silent. Experiments with glycinergic blockade provide independent evidence for inspiratory excitation of post-I neurons. After strychnine, post-I neurons discharged during inspiration in adult mice (Büsselberg et al., 2001) and neonatal rats (Dutschmann and Paton, 2002), and the three-phase rhythm of the mice became two-phase. The inspiratory excitation supported here is a candidate source of that discharge. Together with aug-E-pattern excitation of pre-I/I, it supports excitatory interactions from inspiratory to expiratory populations and from expiratory to inspiratory populations. Populations with separate discharge phases are therefore linked by excitation as well as by inhibition.

### 4.4 Resolution of presynaptic assignments

In our analysis, the overlapping pre-I/I, ramp-I, and early-I activity patterns permitted some individual input assignments while leaving others unresolved. For post-I excitation, ramp-I dominated individual fits, but pre-I/I inputs remained a statistical alternative. For ramp-I excitation, we distinguished a ramp-I contribution and found support for additional inspiratory input. Figure 4 separates supported combined inputs from these finer assignments and alternatives.

Fine assignments depend on differences among representative waveforms. A systematic template mismatch, synaptic filtering, or an unrepresented input can create an apparent need for another pattern across many cells, particularly for early-I-pattern excitation of pre-I/I. Increasing the postsynaptic sample cannot remove shared template bias. Combined inspiratory assignments distinguish broader input phases and are less dependent on these within-phase differences.

### 4.5 Cell identity and anatomical connectivity

The five activity profiles used in our reconstruction come from recordings in three regions of the ventral respiratory column. In the intact pontine-medullary preparation, Smith et al. (2007, their Figure 4A) recorded post-I and aug-E activity in the BötC, pre-I/I and early-I activity in the pre-BötC, and ramp-I activity in the rVRG. Marchenko et al. (2016) also recorded post-I and aug-E population activity together in the BötC in vivo. In adult rats, Sun et al. (1998) classified 80% of the respiratory neurons they recorded in the BötC as expiratory, compared with 22% in the pre-BötC. Our intracellular recordings included all five discharge patterns in the pre-BötC, with additional post-I and aug-E neurons in the BötC (Molkov et al., 2025).

In the circuit proposed by Smith et al. (2007, their Figure 7), expiratory populations in the BötC inhibit the inspiratory network in the pre-BötC. Activating inhibitory neurons in the BötC suppressed pre-I/I discharge in the pre-BötC (Ausborn et al., 2018). This regional effect and the predominance of expiratory neurons in the BötC make it a plausible source of the post-I-pattern inhibition we found in all three inspiratory populations.

We identified glycinergic post-I neurons in both the BötC and the pre-BötC (Molkov et al., 2025). In rhythmic pre-BötC slices, Molkov et al. (2026a) inferred reciprocal inhibition between inspiratory and expiratory populations. Their Figure 8 compares this local inhibitory circuit with the corresponding interactions in our earlier in situ reconstruction. Thus the pre-BötC itself contains inhibitory circuitry capable of coordinating inspiratory and expiratory activity. How much of the post-I-pattern inhibition in situ originates in each region remains unknown.

Even within one region, neurons with similar discharge can differ in transmitter and projection (Smith et al., 2013). Immunolabelling identified inhibitory neurons in each of the five discharge patterns, but too few cells were labelled to compare synaptic inputs by transmitter within each pattern (Molkov et al., 2025). The incoming connections in Figure 4 therefore describe the mean inputs to neurons with a given discharge pattern.

### 4.6 Measurement and statistical limitations

Conductance reconstruction depends on stable current-voltage relations and effective reversal potentials (Molkov et al., 2025). Voltage-dependent currents, electrode effects, and reversal uncertainty can distort the profiles. Reversal errors can transfer temporal structure between excitation and inhibition, particularly affecting small excitatory components beside large inhibition. The contributions describe dynamic conductances on a shared normalized scale.

Provisional cell classifications introduce population-assignment uncertainty, and unavailable preparation identifiers prevent estimation of dependence among cells from the same animal.

Precision varies with sample size, especially for the eight early-I cells. Skewed score distributions make the *t*-based significance levels approximate.

The inferred connections summarize separate omission tests under the fixed waveform representation. Unshown connections remain possible. Whether the inferred circuit interactions suffice to sustain a rhythm requires dynamical testing.

### 4.7 Conclusions

We have shown how a novel functional connectome of medullary respiratory circuits can be reconstructed from a large data set of single cell synaptic input conductances, combined with statistical tests for their sources of presynaptic population activity. A unique feature of our data and analyses is that they provided phase-resolved synaptic inputs throughout the respiratory cycle for five major electrophysiological phenotypes of active respiratory neurons. Our findings support the following interpretation of circuit operation: (1) The inhibitory inputs we identified are consistent with the proposed role of inhibition in respiratory phase switching; decrementing post-I inhibition was the largest fitted expiratory input to each inspiratory population. (2) Synaptic excitation increases during inspiratory and expiratory phase discharge ramps alongside declining inhibition. (3) Inspiratory excitation of silent expiratory populations, and aug-E-pattern excitation of pre-I/I support excitatory interactions across the inspiratory-expiratory division in both directions. (4) Rising aug-E excitation together with pre-I/I recurrent excitation and declining post-I inhibition provides a candidate mechanism for recruitment of the next inspiration. The next step will be to evaluate the dynamical operation of our reconstructed circuit by simulations incorporating the proposed synaptic interactions and neuronal biophysical properties.

## Appendix A: Conductance reconstruction and statistical inference

### A.1 Phase-specific conductance reconstruction

Figure A1 connects intracellular recordings, phase-specific current-voltage relations, and reconstructed conductances. The full reconstruction method is described by Molkov et al. (2025). For the present analysis, the fitted electrical relation at each phase is

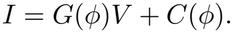

In current clamp, fitting *V* = *a* + *bI* gives *G* = 1*/b* and *C* = *−a/b*. In voltage clamp, we estimate *G* and *C* directly from the current-on-voltage regression. Before minimum subtraction,

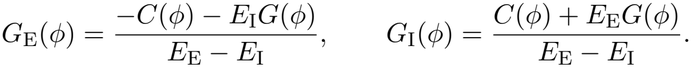

Subtracting each component’s cycle minimum gives its dynamic part. The summed minima define an effective reference conductance; reconstruction required this reference to be positive.

We estimated the inhibitory reversal potential by fitting an upper supporting line to the phasewise (*G, C*) points, minimizing its summed vertical gap. Its slope is *−E*_I_. Under the separation assumption, this boundary corresponds to the minimum excitatory component. We constrained *E*_I_ to [*−*110, *−*60] mV; the estimate reached one of these limits in 58 of 168 recordings. We fixed the effective excitatory reversal potential at *E*_E_ = *−*20 mV. Section 2.5 describes phase alignment and normalization.

### A.2 Normalization and individual conductance fits

For one recording, let *g*_E_ and *g*_I_ denote the dynamic conductances after subtraction of their respective cycle minima. We normalized both profiles by their shared maximum:

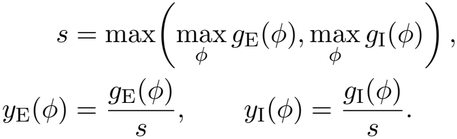

For either conductance type, let *y* contain the 100 phase-bin values of the normalized profile and let the columns of *A* contain the five activity templates. We obtained the individual fit by non-negative least squares:

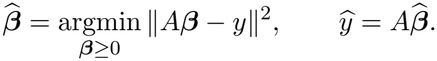

Each coefficient scales one candidate presynaptic contribution. All phase bins entered the objective equally, and no offset was fitted. The recorded conductance is measured above its lowest value in the cycle. If one presynaptic input declines as another rises, both may still contribute at that point, even though the recorded dynamic conductance is zero.

**Figure A1:**
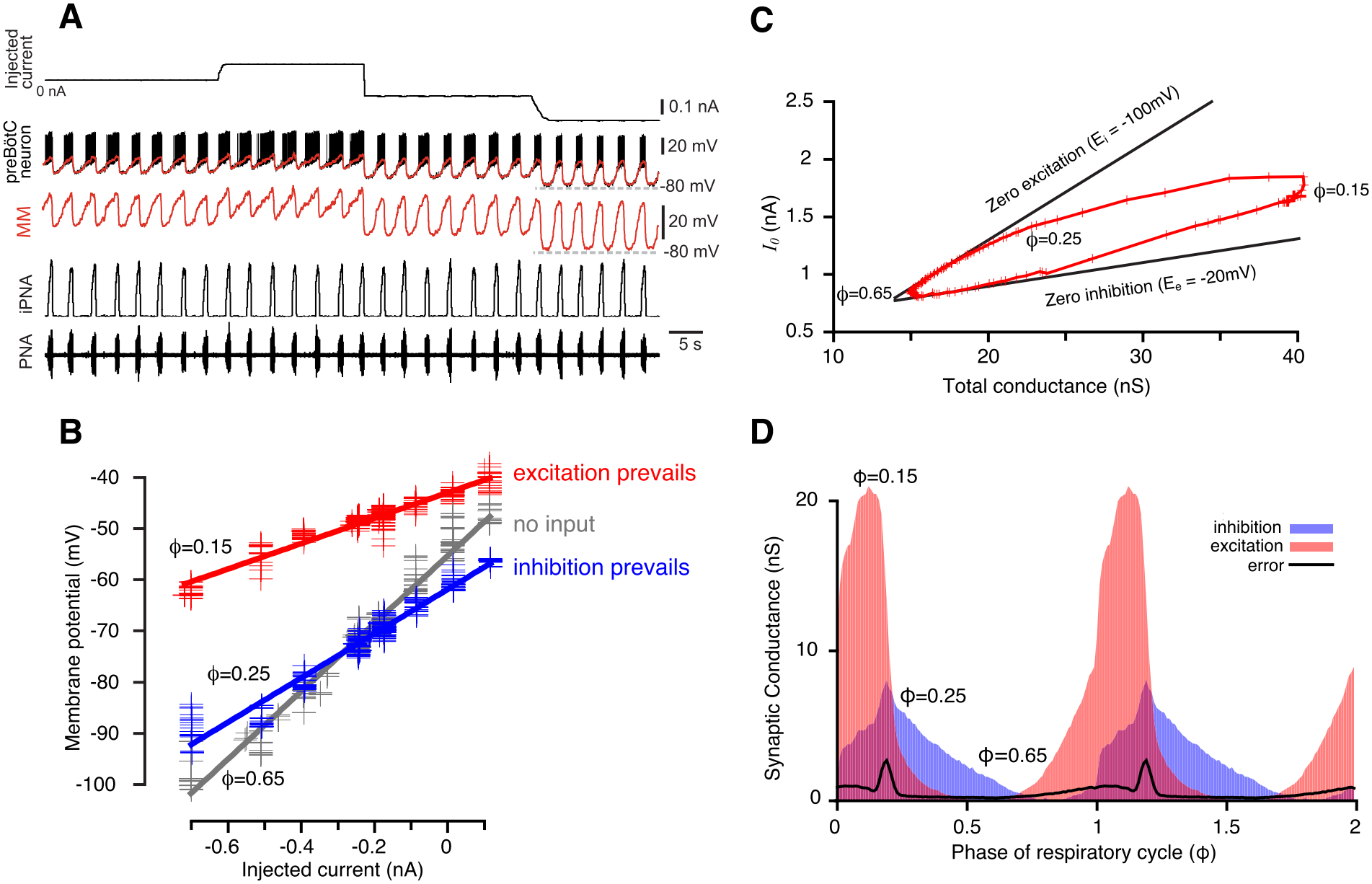
From recordings to conductance profiles. A, current injection and membrane voltage, with moving-median filtering (MM, red), phrenic nerve activity (PNA), and integrated PNA (iPNA). B, current-voltage regressions at three respiratory phases. C, wedge diagram of total conductance and current intercept; upper and lower boundaries represent zero dynamic excitation and inhibition under the original assumptions. D, dynamic excitation (red), inhibition (blue), and reconstruction error (black). Integer phases mark inspiratory onset. Original settings and nS units are retained; present processing is described in Section 2.5. Reproduced from Figure 1 of Molkov et al. (2025), *eLife* 13:RP101959, doi:10.7554/eLife.101959.4, under CC0 1.0.

### A.3 Statistical tests of omitted inputs

We constructed template-derived contrasts and combined one-sided *t* tests with closed testing to assess a nonnegative population-mean representation. The underlying constrained inference problem is established (Fang et al., 2023); the construction below applies it to candidate synaptic inputs.

#### A.3.1 Omission hypothesis and contrast

For one receiving population and conductance type, let *y*_*i*_ be the normalized profile of cell *i*, after averaging recordings within clamp mode and then across modes. Let *A* = [*a*_1_, …, *a*_5_] contain the fixed pre-I/I, ramp-I, early-I, post-I, and aug-E activity templates. We assume

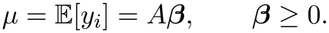

If *D* indexes omitted templates and *K* indexes those retained, the omission hypothesis is

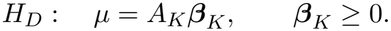

For each omitted template *j ∈ D*, form its NNLS approximation from the retained templates and the residual

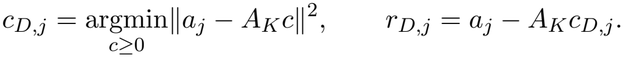

The residual isolates timing features that the retained mixture cannot reproduce. NNLS projection properties (Lawson and Hanson, 1995; Moreau, 1962) give

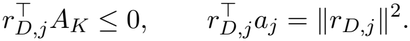

For a nonzero residual, define

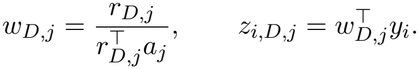

The weights are determined from the templates alone and applied to every cell. Under *H*_*D*_,

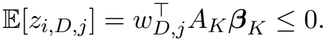

Thus, a significantly positive mean score rejects every nonnegative mixture of the retained patterns. A zero residual means that this contrast cannot distinguish the omitted template.

##### A.3.2 Across-cell test and multiple comparisons

Let *z*_*D,j*_ and *s*_*D,j*_ be the mean and sample standard deviation of the *n* cell scores. We use

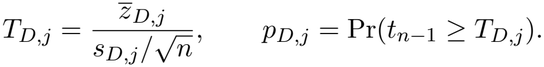

Zero is the largest mean permitted by the omission hypothesis. For independent normally distributed scores with common positive variance, the test is exact at this boundary and conservative for negative null means. The observed score variance incorporates covariance between phases within each cell; the number of cells determines the sample size.

For the inspiratory indices *I* = *{*1, 2, 3*}*, we test all seven nonempty omissions *D ⊆ I*. A local Bonferroni correction combines the contrasts within an omission:

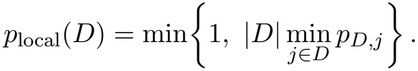

The five-template matrix *A* has full column rank. Under the full mean representation, its coefficients are unique and *H*_*D*_ = _*j∈D*_ *H*_*{j}*_. Closed testing therefore requires rejection of every local omission containing *D* (Marcus et al., 1976):

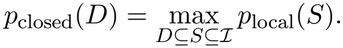

For example, individual ramp-I support requires rejection of the ramp-I omission, both pairs containing ramp-I, and the whole-inspiratory omission. Joint omission can be rejected while individual assignments remain unresolved.

Across five receiving populations and two conductance types, there are ten inspiratory families and twenty single-template expiratory tests. We correct across these thirty families or tests:

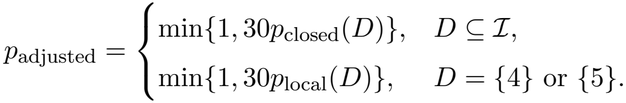

With valid elementary P values, this procedure controls the familywise error rate at 0.05 across all 90 omission hypotheses, allowing dependence between tests. The observed departures from normal score distributions make this control approximate for the present data.

##### A.3.3 From rejected omissions to circuit alternatives

An adjusted P value below 0.05 supports the omitted input or set. Let *R ⊆ I* be a retained inspiratory combination. To be consistent with all rejections, it must satisfy

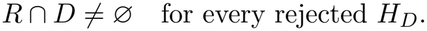

We display the combinations for which no member can be removed while preserving this condition. Any larger combination containing a displayed alternative remains possible. Both expiratory templates remain available.

##### A.3.4 Worked example: inspiratory excitation of post-I neurons

The 54 post-I cells illustrate how an input can be supported while its origin remains unresolved (Figure A2). Omitting all three inspiratory templates leaves only post-I and aug-E. The ramp-I-derived contrast then gives *t*(53) = 9.98 and an adjusted P value of 4.13 *×* 10^*−*12^, supporting an inspiratory excitatory contribution.

Omitting ramp-I alone allows the other inspiratory templates to contribute. The resulting contrast gives *t*(53) = *−*1.72 and an adjusted P value of 1. Joint omission of pre-I/I and ramp-I is rejected, while neither is individually identified. Either can therefore remain as the sole inspiratory template alongside the two expiratory templates; both may contribute. Although ramp-I accounts for 89.5% of the fitted inspiratory excitatory area on average, its larger fitted share does not distinguish it uniquely.

**Figure A2:**
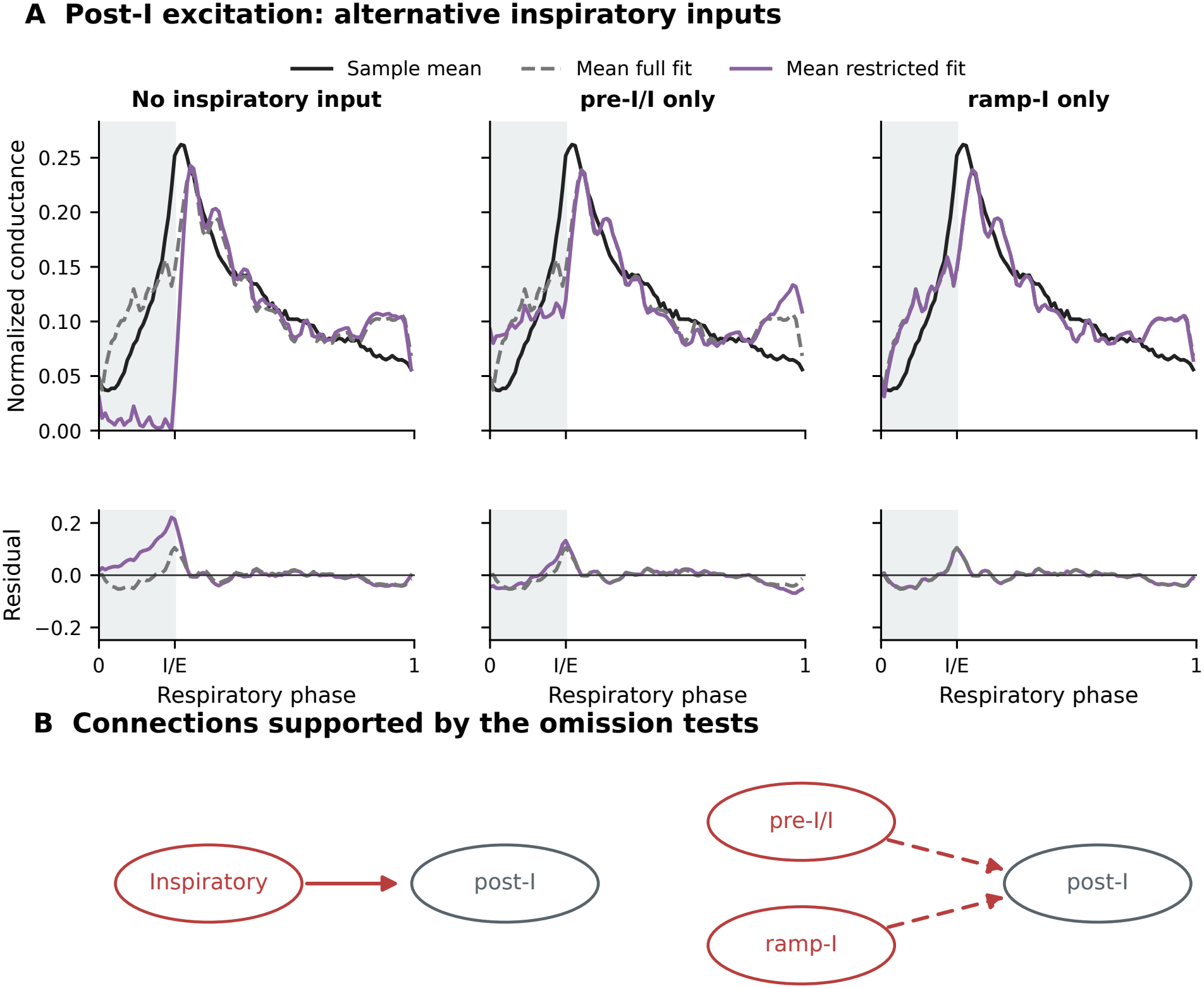
Alternative origins of post-I excitation. A, mean conductance in 54 post-I cells (black), mean individual five-template fits (grey dashed), and mean individual restricted fits (colored). Restricted fits retain no inspiratory template, pre-I/I alone, or ramp-I alone; both expiratory templates remain available. Lower panels show mean conductance minus mean fit. Normalization and respiratory phase follow Figure 2; grey shading marks inspiration. B, omission tests support inspiratory excitation (solid arrow), with pre-I/I and ramp-I as unresolved alternatives (dashed): at least one is needed, and both may contribute. The neutral target pools post-I cells across transmitter identities. Panel A describes fit quality; panel B summarizes across-cell statistical tests.

## Additional information

### Data and code availability

The 168 recordings from 150 cells, activity templates, reconstructed conductances, individual fits and statistical results are archived at https://github.com/ymolkov/resp-net-paper. The repository includes the code and instructions for reproducing the analyses and figures, together with the interactive circuit atlas. Supporting Data S1 provides the fitted contributions, omission-test results and underlying contrasts.

### Author contributions

Y.I.M., I.A.R. and J.C.S. conceived and designed the study. J.C.S. provided the recordings from the Cellular and Systems Neurobiology Section, NINDS, Bethesda, MD, USA. I.A.R. and J.C.S. supervised the work. Y.I.M. developed the analytical and statistical methods, wrote the software and prepared the figures. Y.I.M. and J.C.S. analyzed the data and drafted the manuscript. Y.I.M., I.A.R. and J.C.S. interpreted the results, critically revised the manuscript and acquired funding.

### Generative AI statement

We used OpenAI Codex to help write and test analysis code and prepare figures. We used both Codex and Anthropic Claude to revise the manuscript. We verified the conductance reconstructions and statistical decisions by rerunning the archived code on the recordings. The authors take responsibility for the analyses, their interpretation and the manuscript.

